# Hippocampal theta codes for distances in semantic and temporal spaces

**DOI:** 10.1101/611681

**Authors:** Ethan A. Solomon, Bradley C. Lega, Michael R. Sperling, Michael J. Kahana

## Abstract

The medial temporal lobe (MTL) is known to support episodic memory and spatial navigation, raising the possibility that its true function is to form “cognitive maps” of any kind of information. Studies in humans and animals support the idea that the hippocampal theta rhythm (4-8 Hz) is key to this mapping function, as it has been repeatedly observed during spatial navigation tasks. If episodic memory and spatial navigation are two sides of the same coin, we hypothesized that theta oscillations would also reflect relations between explicitly nonspatial items, such as words. We asked 189 neurosurgical patients to perform a verbal free-recall task, of which 96 had indwelling electrodes placed in the MTL. Subjects were instructed to remember short lists of sequentially-presented nouns. We found that hippocampal theta power and connectivity during item retrieval coded for semantic distances between words, as measured using word2vec-derived subspaces. Additionally, hippocampal theta indexed temporal distances between words after filtering lists on recall performance, to ensure adequate dynamic range in time. Theta effects were only noted for semantic subspaces of 1-dimension, indicating a substantial compression of the possible semantic feature space. These results lend further support to our growing confidence that the MTL forms cognitive maps of arbitrary representational spaces, reconciling longstanding differences between the spatial and episodic memory literatures.

## 1 Introduction

Our ability to form associations rapidly and efficiently has inspired a powerful conceptualization of brain function: It constructs internal “maps” of related information that we mentally navigate when the behavioral context demands it [1, 2, 3]. These maps need not represent only physical layouts, like an office plan or a route to work – cognitive maps could also reflect associations in domains such as autobiographical memories, perceptual inputs, or social interactions [4]. In this way, for instance, we can conceptualize recollection of a life event as taking a mental stroll through a map of our prior experiences, guided by the similarities between items in a multidimensional feature space.

Evidence from studies of memory and spatial navigation provide compelling support for the idea that the brain has a domain-general cognitive mapping ability with a common neural substrate. Decades of lesional, electrophysiological, and imaging studies demonstrate that the human medial temporal lobe (MTL), including the hippocampus and parahippocampal gyrus, support cognitive operations underlying both spatial navigation and episodic memory [5, 6, 7, 8], while studies of rodent navigation have shown that MTL neurons code for spatial locations in a context-dependent manner [9]. More recently, microwire recordings in humans have begun to bridge the human-animal divide, showing the presence of place-coding cells in the MTL whose firing rates can be modulated by task contexts and reflect associations between spatial locations and behaviorally-relevant content [10, 11, 12].

Common to studies of spatial navigation in humans and animals is the emergence of a low-frequency oscillation in the hippocampal local field potential (LFP), called the theta rhythm (typically 4-8 Hz). In rodents, the spiking of place-coding cells occur at drifting phases of the theta rhythm as an animal explores its environment [13, 14], and these theta-modulated firing patterns are re-instantiated as an animal retrieves information about a previously-explored layout [15]. Though such instances of “phase precession” have not been observed in navigating humans, other studies strongly support the general notion of theta phase coding as humans perform spatial and memory tasks [16, 17, 18]. Independent of their relation to neuronal spiking, theta oscillations have been consistently observed in humans during navigation, and tasks that require the encoding or retrieval of spatial information [19, 20, 21, 22, 23].

Though it is now clear that theta oscillations correlate with the formation of spatial relations – or maps – there is a more tenuous link between theta and explicitly nonspatial memory. If the MTL truly serves a domain-general mapping function that creates relational networks between information of all types – be it spatial, semantic, autobiographical, or otherwise – we should expect to observe the same theta-mediated phenomena during nonspatial memory processing. Indeed, theta activity has been observed in human MTL and neocortex during episodic memory processing for words and pictures [24, 25, 26, 27]. However, these studies examine theta through the broad lens of successful-versus-unsuccessful memory, called the subsequent memory effect (SME). Though a useful paradigm, the SME is not a direct test that theta specifically supports the formation of associations between items in memory, i.e. a cognitive map of recently-acquired information. Moreover, many similar studies report the opposite effect: a hippocampal or cortical decrease in theta power during successful encoding or retrieval [28, 29, 30, 31, 32, 33]. These conflicting findings raise serious questions about whether the neural processing of spatial and episodic information rely on the same theta-based mechanisms.

We set out to directly ask whether the established neural signature of spatial memory and navigation – MTL theta oscillations – are also present during the retrieval of relational verbal memory. To do this, we tested 189 human neurosurgical patients with indwelling electrodes on a verbal free-recall task. Prior studies indicate that humans principally rely on temporal and semantic relatedness to organize verbal information [34, 35], so our overarching goal was to ascertain whether human behavior and neural activity aligned with measures of temporal and semantic association. We explicitly tested whether theta rhythms in the human MTL code for (1) semantic distances between retrieved nouns and (2) temporal distances based on the serial position of presented items during encoding. In doing so, we sought to establish whether episodic memory is analogous to spatial navigation. Is the encoding of word items akin to guiding a subject through an abstract space, and is retrieval akin to asking them to walk through it themselves?

Using a pretrained word2vec [36] representation of words in a vector space, we found that we could predict a subject’s recall behavior based on inter-word distances in word2vec subspaces of varying dimensionality. Next, we asked whether inter-word distances, and temporal distances, correlated with pre-retrieval theta power in the hippocampus and parahippocampal gyrus (PHG). We discovered that pre-retrieval hippocampal theta power significantly predicts inter-word distances in semantic space, but theta was not correlated with temporal distances. However, analytic corrections for the case of temporally-restricted recalls caused a time-associated theta effect to emerge. Finally, we assessed inter-regional theta connectivity, finding that hippocampal-PHG coupling also predicted inter-word semantic distances. Taken together, we established that retrieval-associated theta activity encodes representational distances in a word space, supporting the idea that theta underlies the formation of domain-general cognitive maps in the MTL.

## 2 Results

### Clustering effects in recall

We first sought to examine the degree to which distances between items’ semantic representations predicted both recall behavior and neural activity. Neurosurgical patients with indwelling electrodes (N=189) studied and subsequently recalled 12-item word lists. During the study phase, each word appeared for 1.6 seconds followed by a 800-1200 ms blank interstimulus interval (see Fig. 1A for an example list). Following list presentation, subjects first performed a 20-second arithmetic distractor task and then attempted to freely recall the items from preceding list in any order. Subjects completed multiple sessions of this free-recall task, with each session comprising 12-25 study-test lists.

**Figure 1:**
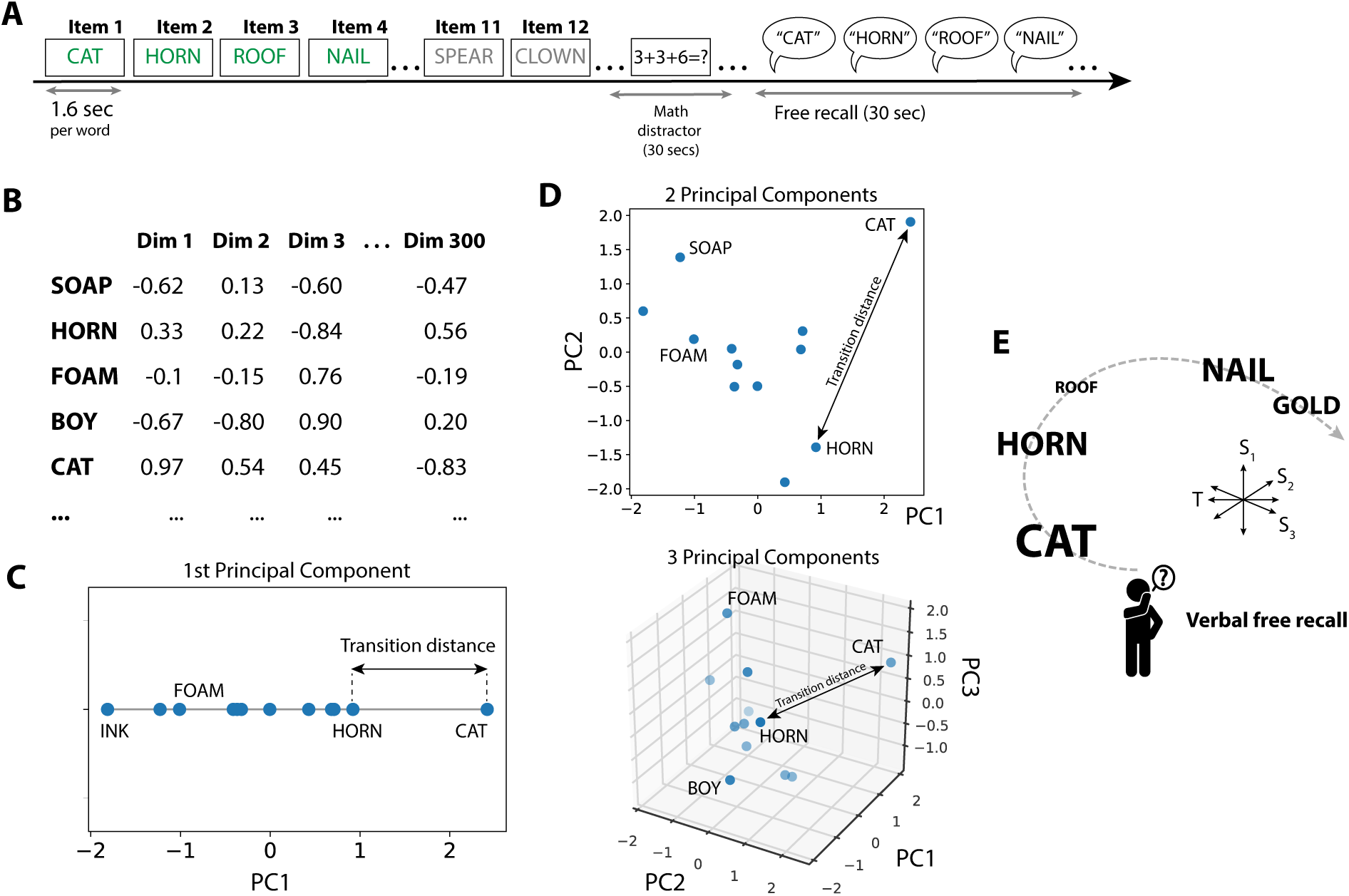
Embedding of word items in representational semantic spaces. **A.** 189 subjects performed a verbal free-recall task, in which they were asked to remember 12-item lists of simple nouns. After a distractor, subjects freely recall as many words as possible from the prior list, which can be conceptualized as a cognitive transition from one word to another. Successful recalls are indicated in green for an example list. **B.** Each word occupies a position in the pretrained 300-dimensional word2vec space (see Methods). PCA is used to extract the principal dimensions of variation for each word list. **C.** Example inter-recall distance in a 1-D semantic space, the first principal component. **D.** The 2-D and 3-D embedding of the words from this list. **E.** Navigation through a 2-D spatial layout is associated with hippocampal theta oscillations, and can be considered analogous to episodic retrieval processes through a multidimensional semantic-temporal space.

We next used a word2vec semantic representation to create layouts of the words in each list, at varying dimensionalities. Word2vec is a method for creating vector representations of words based on regularities present in text corpi [36]. Google has made available a pre-trained set of 300-dimensional word vectors based on a large Google News text corpus. Word2Vec makes explicit the multidimensional nature of word representations, enabling us to ask whether the dimensionality of a word subspace is related to behavioral or neural variables of interest. Accordingly, we used principal component analysis (PCA) to select the top *n* orthogonal dimensions which explain the greatest variance among list words in their native 300-D word2vec space (1B-D; see Methods). The result, for each list, is a set of coordinates that locate each word in an *n*-dimensional space, where *n* is a maximum of 11 (since each list consists of 12 words).

Do distances in these semantic representational spaces align with how subjects retrieve words during the free-recall phase? Prior literature has established that other measures of semantic similarity predict the order of word recall, in that semantically-related words will tend to be retrieved in clusters [34]. Our expectation was that distances in these word2vec spaces would also predict clustered recalls, and they did: Using the semantic clustering score – a metric which indexes whether a subject tends to organize recalls according to semantic relatedness – subjects exhibited recall patterns which were significantly correlated with distances in reduced word2vec spaces of any dimensionality (Fig. 2A, Fig. 2B; 1-D clustering *t*(188) = 7.4, *P*<0.001). Semantic clustering scores tended to rise with increasing PCA dimensionality, but beyond two dimensions, this rise did not significantly outpace the increase in overall explained variance with additional dimensions; regressing out explained variance from the slope of the curve in Fig. 2B showed, across subjects, a significant (*P*<0.05) rise between one and two dimensions (Fig. 2A inset; Fig. 2C; see Methods).

**Figure 2:**
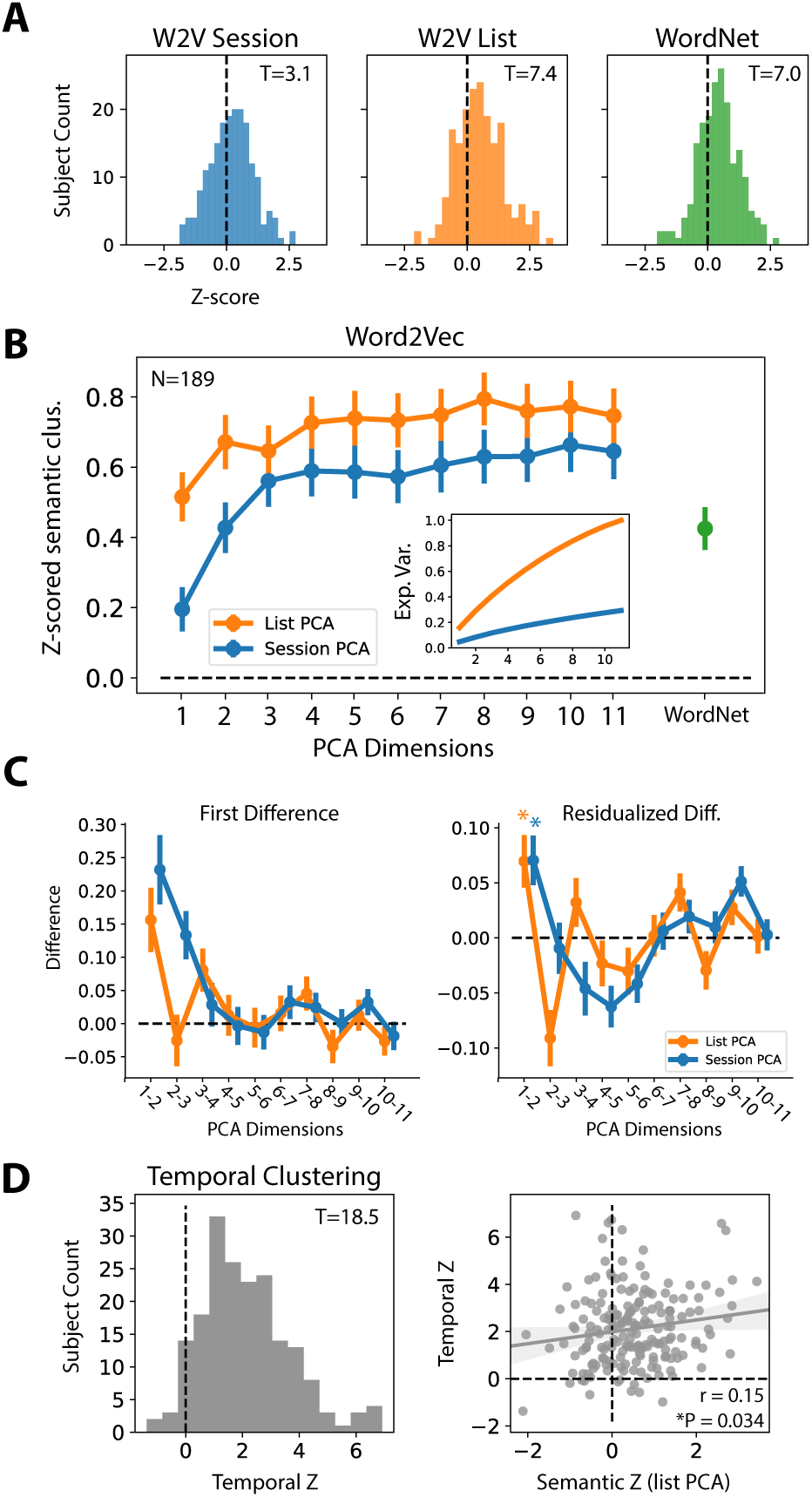
Recall patterns are predicted by inter-word distances in semantic spaces. **A.** Assessing whether inter-word distances in 1-D word2vec-derived spaces predicted the recall of semantically-related words. For each subject, a semantic clustering z-score was computed, reflecting the degree to which their pattern of recalls correlated with inter-word distances in PCA-reduced word2vec spaces of *n*-dimension (see Methods), here shown for just the 1-D space reflecting the first principal component. PCA was applied to the 12-vector group of words from each list individually (list-PCA; orange), or applied to the vectors of all words seen in an experimental session together (session-PCA; blue). As a benchmark, WordNet lexical similarities were also used to calculate semantic clustering (green). Population-level semantic clustering was significant for all methods (1-sample *t*-test, *P*<0.002). **B.** Analysis from (A), extended to PCA-derived semantic subspaces of higher dimensions. *Inset:* Cumulative explained variance for list- and session-level PCA. *Right:* Mean semantic clustering score using WordNet-derived semantic relations. **C.** *Left:* First-order differences of the clustering vs. dimensionality curves in (B). *Right:* First-order differences residualized on the explained variance, indicating that behavioral clustering increased at a faster rate than the explained variance between the first and second dimension for both list-PCA and session-PCA (*P*<0.05). **D.** *Left:* Histogram of subjects’ temporal clustering scores, which were significantly above chance (*P*<0.001). *Right:* Correlation of subjects’ semantic clustering and temporal clustering scores, indicating a significant positive relationship (*r* = 0.15, *P*=0.034).

To ask whether semantically-clustered recalls also correlated with inter-word distances in other spaces, we examined the relationship between sequential recalls and reduced word2vec spaces derived from PCA on all of the words presented in an experimental session (typically 144-300 words, depending on the number of lists done). In this way, PCA is identifying the orthogonal dimensions that explain the greatest variance for all the words seen over the course of the session, as opposed to words seen within a list. Semantic relatedness from session-level PCA also predicted clustering during free recall for all dimensions, though to a lesser degree than list-level PCA (Fig. 2A, Fig. 2B; 1-D clustering *t*(188) = 3.1, *P*=0.002). Clustered recalls were better predicted by inter-word distances at higher dimensions – like we found in list-level PCA – but as before the rise in clustering scores did not outpace the increase in explained variance beyond two dimensions (Fig. 2A inset, Fig. 2C). As a benchmark, we also showed that recalls clustered according to inter-item similarities derived from the WordNet lexical database [37], which does not rely on a model-based embedding of words in a high dimensional vector space (Fig. 2A, right; Fig. 2B, right; 1-sample *t*-test, *t*(188) = 7.0, *P*<0.001).

These data replicate prior findings that verbal recall is governed in part by semantic relations [34], using a multidimensional framework for capturing inter-word similarities. However, it is well-known that humans also organize verbal memory according to the temporal sequence of items during initial encoding [35]. The temporal clustering score quantifies this relationship by assessing the likelihood that a subject will recall two items in sequence, given their serial position during encoding. As in prior studies, we showed a strong population-level effect of temporal clustering (1-sample *t*-test, *t*(188) = 18.5, *P*<0.001). To ask whether the tendency to semantically cluster is related to the tendency to temporally cluster, we found that a subject’s temporal clustering score was weakly but significantly correlated with their 1-D list-PCA semantic clustering score (*r* = 0.15, *P*=0.034).

Taken together, we demonstrated that humans behaviorally cluster their recalls of verbal items according to inter-word distances in word2vec semantic spaces of varying dimension. Altogether, this is not surprising – word2vec representations should not be expected to differ dramatically from other methods of capturing semantic similarity, like latent semantic analysis [38]. In both list-PCA and session-PCA designs, we found that the relationship between recall structure and semantic dimensionality saturated after two dimensions, suggesting that the brain’s organization of semantic features in memory relies substantially (though not exclusively) on one or two principle axes of variation. We next sought to find the neural underpinnings of semantically and temporally-clustered recalls.

### Oscillatory power during search through 1-D spaces

Rodent studies suggest a fundamental link between hippocampal theta oscillations and the organization of sequential information: Neuronal firing becomes associated with a precessing theta phase during exploration of a space, allowing a time-compressed version of sequential items to be encoded within a single theta cycle. Upon reaching a decision point in a spatial layout, it has also been demonstrated that neuronal firing replays along a theta oscillation, akin to the animal retrieving information about places nearby in a previously-explored environment (Fig. 1E). Here we ask: Does theta also support the retrieval of nearby items in a semantic representational space?

To assess this, we recorded local field potentials (LFPs) from depth electrodes in the hippocampus or parahippocampal gyrus (PHG) of human neurosurgical patients. Our general approach was to use the framework outlined earlier (see Fig. 1) to assess whether theta power leading up to a retrieval event predicted the semantic distance between the two subsequently recalled words. A positive correlation between theta power and semantic distance would suggest that theta oscillations are serving to reinstantiate the semantic context established during encoding – much as rodent theta reinstantiates the activity of place cells – and enable a participant to select the optimal path through a representational space (Fig. 1E).

Figure 3 demonstrates this procedure in an example subject. In one list, the subject recalled five of the original 12 words, recalled in an order that sometimes reflects close semantic relations (e.g. recall of “cloud” was followed by “star”), and other times weak semantic relations (e.g. “star” was followed by “nose”; Fig. 3A). For each recall transition, the inter-word distance is captured by the semantic clustering score, here reported as a percentile rank with 1 indicating the closest-possible semantic transition (see Methods). We first computed semantic clustering scores based on the 1-D list-PCA space (and later generalized to higher dimensions). We next measured the theta power in 1-second intervals leading up to each recall event, just prior to vocalization (Fig. 3B). A ∼4Hz theta oscillation is seen preceding the recall of “nose,” which was followed by the recall of “peach” (0.66 clustering score). Finally, we measured the (z-scored) pre-retrieval theta power for all words recalled in an experimental session, and correlated it with the semantic distance between the two subsequently recalled words. This subject shows a significant positive relationship between 1-D semantic distance and pre-retrieval theta power (Fig. 3C, Pearson’s *r* = 0.52, *P*=0.004). To assess the actual degree of theta power change relative to average, we also binned semantic distances into “short” (>0.75 clustering score) and “long” (<0.25), and averaged theta power for recalls in both bins (Fig. 3D). This demonstrated that theta power increased above average prior to near transitions, and fell below average prior to far transitions.

**Figure 3:**
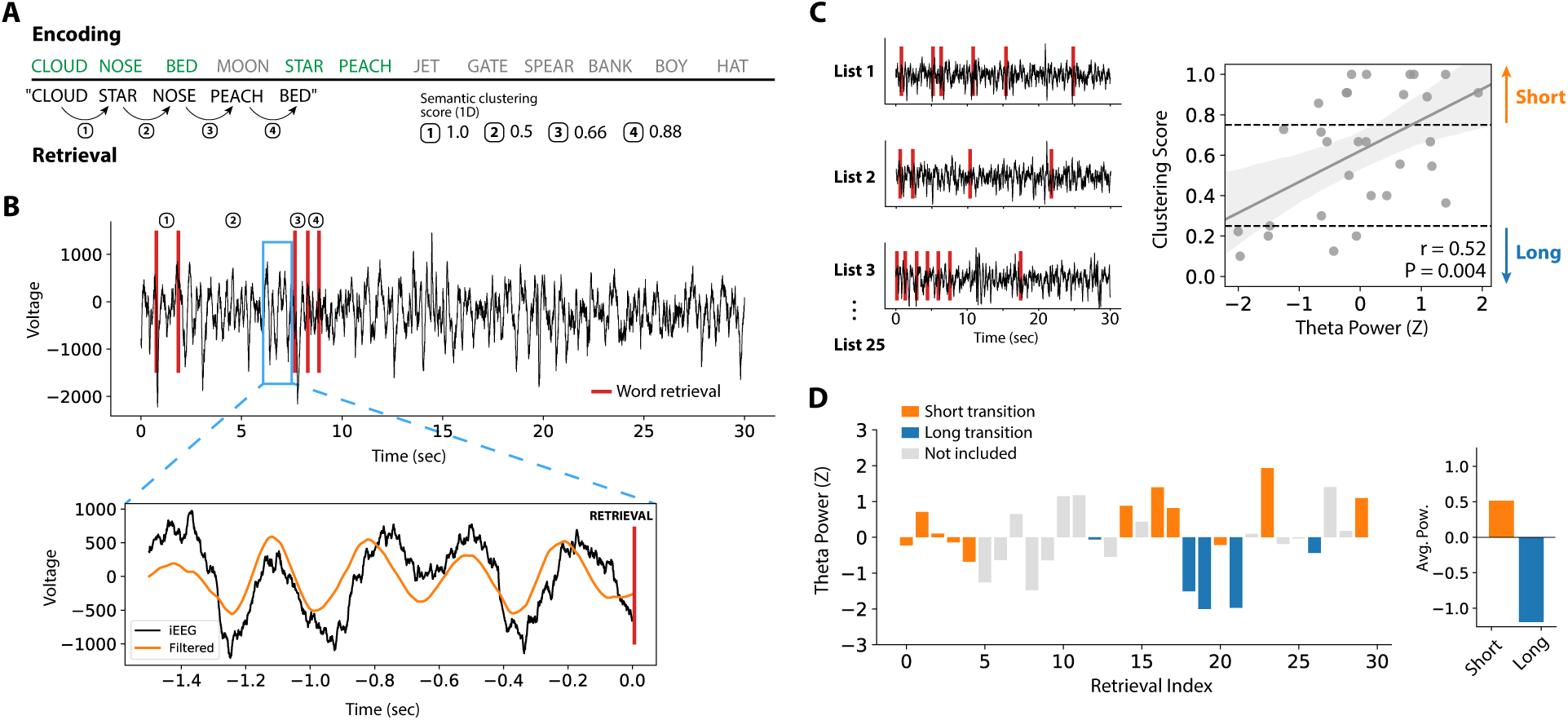
Correlating pre-retrieval theta power with semantic transition distance. **A.** In an example list, a subject recalls five of the original 12 words, making close and far transitions through a 1-D semantic space. Transition distances are measured by a percentile rank called the clustering score, with 1 indicating the closest-possible transition through a given semantic space (see Methods). **B.** Theta power was extracted in 1-second intervals immediately prior to each retrieval event. A ∼4Hz theta oscillation preceding the third recall is highlighted, here measured from an electrode in the CA1 subfield of the hippocampus. **C.** Theta power is measured prior to all retrieval events across an experimental session and z-scored. Z-scored theta power was correlated with semantic transition distance (i.e. the clustering score), demonstrating a significant positive relationship for this subject (*r*=0.52, *P*=0.004). **D.** To further quantify the magnitude of theta power change, inter-word distances were binned into “short” (>0.75 clustering score) and “long” (<0.25), and z-scored theta power was averaged for all retrieval events that fell in each bin. In this subject, theta power increased above average for close transitions, and fell below average for far transitions.

Across all subjects with electrodes placed in the hippocampus (either hemisphere; *N*=70), there was a significant correlation between pre-retrieval theta power and 1-D transition distance (1-sample *t*-test, *t*(69)=2.16, *P*=0.034; Fig. 4A, top; see also Fig. S1), though this effect was primarily driven by activity in the left hippocampus (Fig. S2). Given substantial prior evidence from the literature, we expanded the scope of our analysis to test power in two additional frequency bands: gamma (30-55 Hz), and high-frequency broadband (HFB; 70-150 Hz). Findings in humans and animals implicates MTL gamma oscillations in memory processing [39, 40, 41], and HFB has been used as a general marker of neural activation [42]. In neither band did we find a significant correlation between pre-retrieval power and subsequent transition distance (1-sample *t*-test, *P*>0.05).

**Figure 4:**
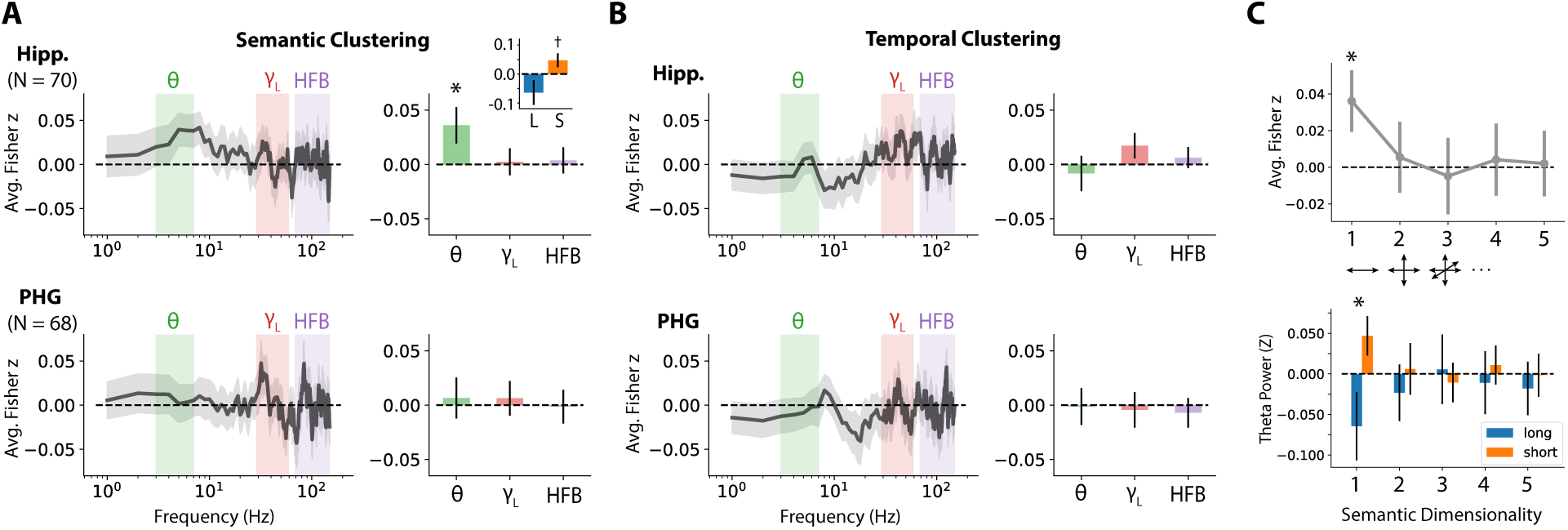
Neural correlates of memory search through 1-D representational spaces. **A.** Correlation between pre-retrieval power and ensuing transition distance in a 1-D semantic space (see Fig. 3 and Methods for details). We assess power-distance correlations in three frequency bands of interest: theta (4-8 Hz), low gamma (30-55 Hz), and high-frequency broadband (HFB; 70-150 Hz). The entire correlation spectrum is shown for completeness, though other bands were not assessed. Band-averaged effects are shown in the bar plot to the right. *Top row:* Power-distance correlations in the hippocampus (N=70 subjects). *Inset:* Average z-scored theta power, binned according “long” and “short” transition distances. *Bottom row:* Power-distance correlations in the PHG (N=68 subjects). **B.** Correlation between pre-retrieval power and ensuing transition distance in a 1-D temporal space, reflecting the serial position of items during encoding. Otherwise structured as in (A). **C.** *Top*: Hippocampal theta predicts semantic transitions only in 1-D semantic space. Pre-retrieval hippocampal theta was correlated with subsequent semantic transition distances in PCA-derived subspaces of varying dimensionality. Only distances in 1-D spaces showed a reliable correlation with theta power. *Bottom:* Similar to above, demonstrating the relative change in theta power for long or short transition distances in subspaces of varying dimensionality. \**P*<0.05, †*P*<0.1. Error bars show +/- 1 SEM. See also Figure S1.

A recent study by Herweg, et al. (2018) noted that theta power in the parahippocampal gyrus (PHG) predicted the recall of items that occurred nearby in a spatial layout [19]. We therefore asked whether PHG theta power demonstrated the same relationship with inter-word transition distances. Unlike our finding in the hippocampus, PHG theta power was not correlated with ensuing transition distances in a 1-D space (1-sample *t*-test, *t*(67)=0.35, *P*=0.73), nor did we find a significant relationship in our two higher frequency bands. Given the strong behavioral tendency for subjects to recall words in a temporally organized manner, we next asked whether distances in a temporal “space” correlated with pre-retrieval power. To do this, we applied the same analysis procedure as for semantic distances, but instead used a measure of temporal transition distance instead of semantic transition distance, i.e. if a subject makes a recall transition between two contiguous words from the original list, the clustering score is 1. We found no significant relationship between pre-retrieval power and temporal transition distance, in any frequency band or region (Fig. 4B, 1-sample *t*-test, *P*>0.05).

By correlating transition distances in 1-D semantic and temporal spaces, we demonstrated that (1) the degree of hippocampal theta power prior to a retrieval event codes for the ensuing semantic transition distance, suggesting that theta serves to reinstantiate semantic context, and (2) short transitions in the time dimension are not significantly predicted by neural activity in our prespecified bands. These findings align with the general idea that theta oscillations support the encoding and retrieval of information organized in a cognitive map. However, it is unclear why the time dimension – shown to powerfully correlate with retrieval behavior – does not have a strong neural correlate (see “Reconciling the absence of time-related theta activity”).

### Multidimensional semantic spaces

We next asked whether distances in higher-dimensional semantic spaces were also predicted by pre-retrieval theta power. Asking this question lets us assess how the brain compresses semantic information – for example, does theta power also correlate with inter-item distances in a two- or three-dimensional semantic layout? Our use of the word2vec semantic representation enabled us to directly test this hypothesis.

Given our prior finding that pre-retrieval theta predicts 1-D semantic distances, we extended our analysis to ask whether pre-retrieval theta predicts semantic distances in higher dimensions. We found that theta power was not correlated with semantic distances beyond the 1-D subspace (Fig. 4D, 4E; *P*>0.05). Dimensionalities between 1 and 5 are depicted in Fig. 4C; dimensionalities between 6 and 10 also showed no effect (all *P*>0.37). This suggests that theta power as a neural correlate of semantic distances is sensitive to the most-compressed subspace possible, i.e. the first principal component of variation in a list.

Though counterintuitive, this finding is in accordance with our earlier behavioral analyses that showed additional PCA components beyond two dimensions did not add any predictive capacity (see first-difference curves in Fig. 2C). PCA components 3-10 did not correlate with semantic clustering any more than would be expected by their marginal explained variance. In other words, the marginal benefit of additional PCA dimensions beyond the first two was small; accordingly, it is not surprising that theta power is only significantly correlated with distances derived from the first and most behaviorally-relevant component.

### Theta connectivity effects in the MTL

While MTL theta power is a clear correlate of human spatial navigation – and possibly semantic-temporal navigation as well – several prior studies also suggest that memory and navigation are supported by inter-regional theta connectivity. In particular, theta connectivity between the PHG and neocortex has been demonstrated during retrieval of spatial information [43], and between hippocampus and PHG during the encoding or retrieval of word items [44, 31, 19]. More broadly, inter-regional theta connectivity has been hypothesized to facilitate synaptic plasticity and gate the flow of information between brain regions, by providing a consistent phase reference across regions that process different aspects of an animal’s experience [45, 46, 47]. Having shown that recall behavior and local theta power correlate with distances in 1-2 dimensional semantic spaces, we next set out to ask whether inter-regional theta connectivity also correlated with the retrieval of semantic or temporal context.

Figure 5 outlines our procedure in an example subject. We using the phase-locking value (PLV) [48], a measure of the consistency of phase differences between two electrodes, to assess the degree of theta connectivity between PHG and hippocampus. Here, theta-frequency phases are extracted from 1-second intervals prior to the retrieval of each word, for an electrode in CA1 and an electrode in perirhinal cortex (PRC) (Fig. 5A-B). The PLV is computed on the phase differences between the two regions, prior to each retrieval. Figure 5C illustrators how CA1-PRC theta phases are highly consistent prior to a short semantic transition (1.0 clustering score), since the theta peaks in CA1 always correspond to the falling phase of the oscillation in PRC. Prior to a far semantic transition (0.1 clustering score), the theta peaks in CA1 do not align consistently with the ongoing PRC oscillation. We also note that, in order to avoid assessing phase-locking between bipolar pairs that span the hippocampus and PHG but share a common monopolar contact, our connectivity analyses utilized an MTL-average reference (see Methods for details).

**Figure 5:**
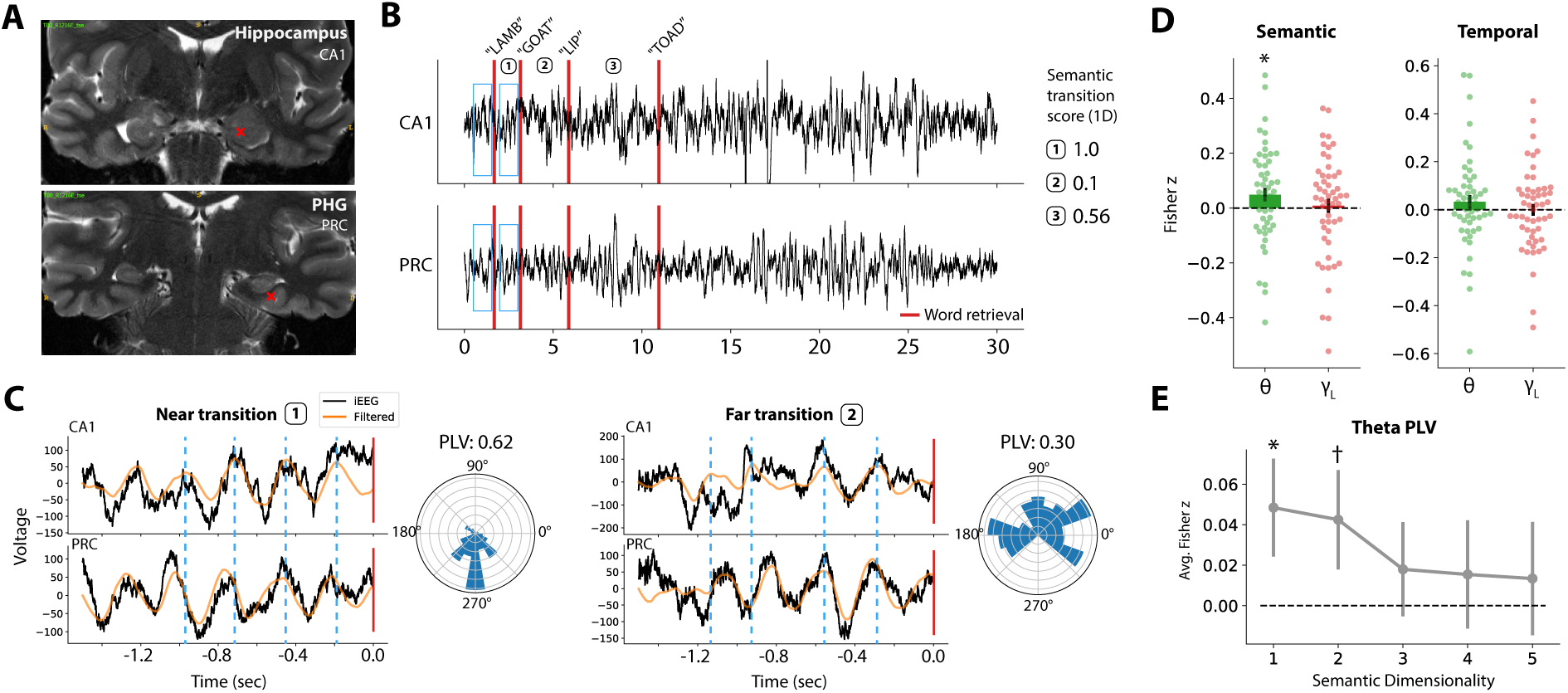
Assessing pre-retrieval theta coupling between the PHG and hippocampus. **A.** In an example subject, phase-locking was assessed between bipolar contacts localized to the hippocampus (CA1) and PHG (perirhinal cortex; PRC), indicated as red crosses. **B.** As in the power analyses, LFP is extracted from each electrode, and theta coupling is assessed in the 1-second intervals leading up to vocalized recalls. A score is assigned to each transition, capturing the closeness of successive recalls in semantic or temporal dimensions. **C.** Phase locking is measured by assessing the consistency of theta phase between the electrodes, for each transition. A close semantic transition is depicted on the left, where the peak of the CA1 oscillation (dashed lines) consistently aligns with the falling phase of the PRC oscillation. A far semantic transition is shown on the right, with inconsistent phase alignment. The concentration of phase differences over time is quantified as the phase-locking value (PLV), visualized as polar histograms. For the example trials shown, the close semantic transition has a higher PLV (0.62) than the far transition (0.30). **D.** Hipp-PHG theta coupling mediates semantic search. *Left:* Hippocampal-PHG theta coupling significantly correlated with ensuing semantic transition distance in a 1-D space (1-sample *t*-test, *t*(55)=2.00, *P*=0.050). No effect was observed for phase-locking in the gamma band. *Right:* Phase-locking was not predictive of temporal transition distance in either frequency band (*P*>0.05). **E.** Theta phase-locking was marginally significantly correlated with 2-D semantic transition (*t*(55=1.73, *P*=0.089, in addition to 1-D transitions, as noted in (D). \**P*<0.05, †*P*<0.1. Error bars show +/- 1 SEM.

Hippocampal-PHG theta phase locking was measured for all subjects with electrodes in both areas (N=56), and as earlier, was correlated with the subsequent semantic or temporal transition distance. Given the possibility of intra-MTL gamma connectivity [49, 50], we further included that band in this analysis. Across all subjects, we found a significant correlation between hippocampal-PHG theta phase locking and 1-D semantic transition distance (1-sample *t*-test, *t*(55)=2.00, *P*=0.050; Fig. 5D), and no effect in the gamma band (*t*=0.32, *P*=0.75). Extending the analysis to higher semantic dimensions, we found a marginally significant correlation between theta coupling and semantic distances in a 2-D space (*t*(55)=1.73, *P*=0.089; Fig. 5E), but no others. Results were qualitatively similar when regressing out power from PLV values, to correct for possible signal-to-noise changes (Figure S3). Taken together, these data suggest that theta coupling between the PHG and hippocampus mediates successful search through a one- and possibly two-dimensional semantic space.

Does pre-retrieval hippocampal-PHG theta coupling support the reinstatement of temporal context? Theta coupling was only weakly correlated with temporal transition distance, but did not reach statistical threshold (1-sample *t*-test, *t*(55)=1.20, *P*=0.23; Fig. 5D). This result mirrored our earlier finding that local theta power was not correlated with temporal transition distance, even though time was the most prominent organizing factor in recall behavior.

### Reconciling the absence of time-related theta activity

Our finding that MTL theta power and connectivity do not relate to temporal clustering came as a surprise – temporal relations are the most prominent organizing principle of verbal free-recall, and recent studies support the idea that neurons in the MTL encode aspects of time [51]. If time is a key dimension along which humans organize episodic memories, and theta rhythms are the physiological substrate of episodic associations, why are temporally-clustered recalls not associated with increases in MTL theta activity?

We hypothesized that our temporal theta-correlation analysis was affected by any variables that could limit the dynamic range of contextual coding during recall. For example, subjects often exclusively retrieve the first few list items as a group, suggesting that these words are associated with a general list context and that such lists do not reflect a dynamic range of temporal contexts. To account for this possibility, we applied a simple modification to the temporal transition analysis presented in Fig. 4, noting that lists with a small number of words recalled (i.e. four or less) are more likely to be confounded by a lack of temporal dynamic range, while high-performance lists with many recalls are more likely to capture meaningful variability across temporal lags (or “distances”). We therefore computed the correlation between theta power and temporal transition distance at varying list-performance thresholds, and compared our findings to the correlations based on the full dataset (we note that, in all analyses in this manuscript, lists with three or fewer recalls were excluded). Additionally, to ensure that we captured a range of temporal lags, we binned clustering scores into “long” and “short” and directly compared the two groups, as was demonstrated in prior analyses (see Fig. 3D and Fig. 4E).

At higher performance thresholds, we noted a shift towards higher correlations between pre-retrieval theta power and temporal transition distance (Fig. 6). In the 22 subjects with sufficient data in high-performance lists (7 or more recalls per list), higher theta power was significantly associated with shorter temporal transition distances (1-sample *t*-test, *t*(21)=2.17, *P*=0.041; Fig. 6), though the association was not apparent at lower performance thresholds (4-recall threshold; *t*(21)=0.77, *P*=0.44). On a per-subject basis, filtering for list performance yielded a marginally significant difference between the theta effect in all-list versus high-performance conditions (paired *t*-test, *t*(21)=1.79, *P*=0.087). As was demonstrated for semantic transitions, the correlation spectrum revealed a qualitative peak in the theta band (Fig. 6, bottom row).

**Figure 6:**
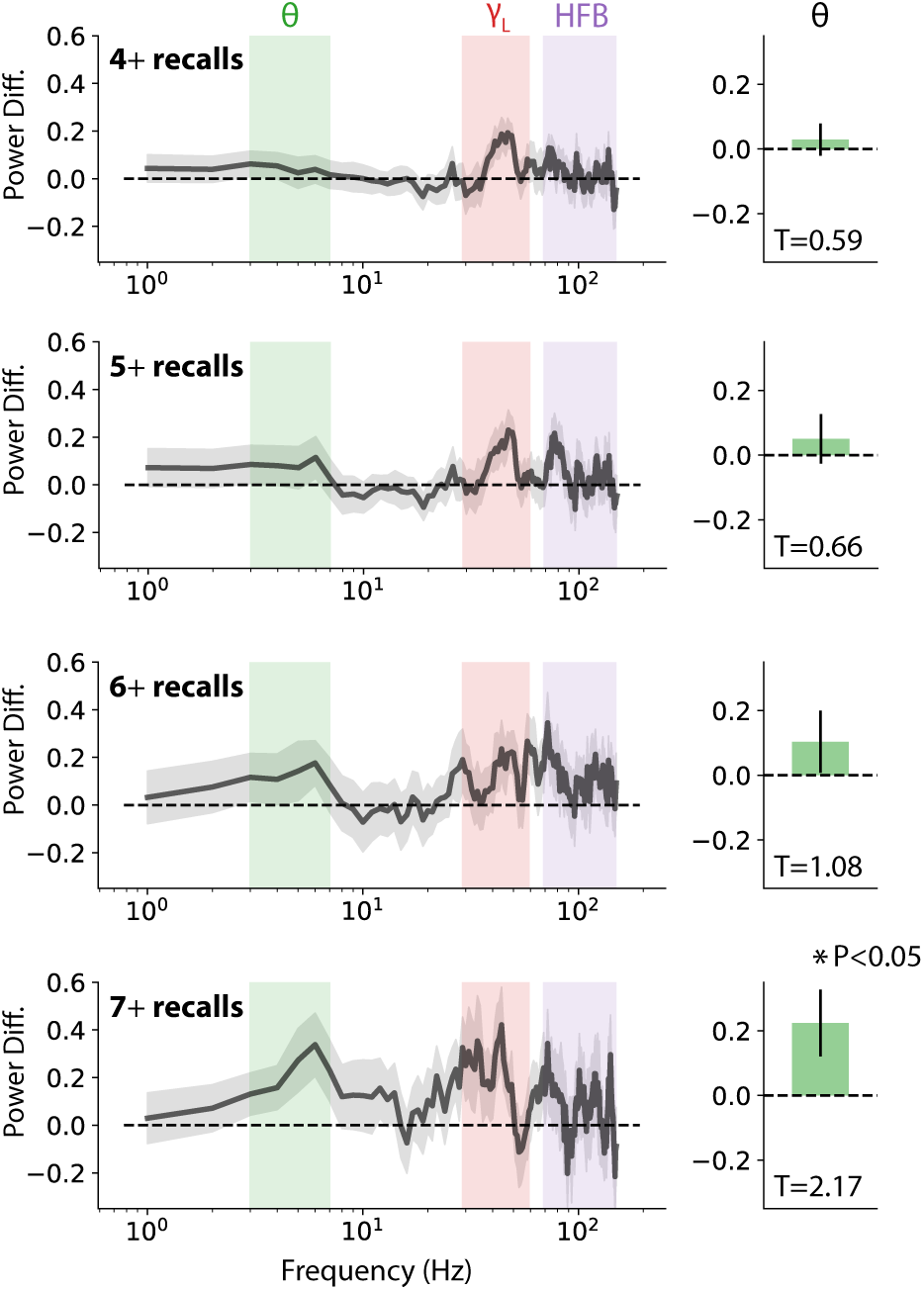
Reconciling the absence of time-associated hippocampal theta. To ask whether restricted temporal dynamic range was contaminating measures of time-associated theta power, recall periods were filtered according to the number of recalled items per list. The association between theta power and temporal transition distance is shown for four different list-performance thresholds, among the 22 subjects with sufficient data at the highest threshold. The theta power difference between short and long temporal transitions (see Fig. 3D) is significantly greater than zero for high-performance lists (7-recall threshold; *t*(21)=2.17, *P*=0.041), and increases monotonically at each threshold. The power difference spectra, reflecting the degree to which theta power is associated with short vs. long temporal transitions, reveals an emerging spectral peak in the theta band, indicating a significant association between theta power and temporal transition distance. \**P*<0.05, error bars show +/- 1 SEM.

By focusing on lists with a high number of recalls, and thereby mitigating potential confounds in our temporal correlation analysis, we found evidence for our hypothesized relationship between hippocampal theta and temporal transition distance. This result mirrored our earlier finding of hippocampal theta associated with close semantic transitions, and since word semantics were unrelated to list structure, no corrections were needed to observe the effect. We note that 22 subjects had sufficient high-performance lists to enter this analysis, relative to the 70 subjects included in the semantic analysis. This introduces a potential confound of a subject’s general memory performance or task ability, and weakens our statistical power. Nonetheless, a performance filter demonstrated the strong possibility that an underlying temporal-theta effect was present, but not directly observable, in our earlier analyses.

## 3 Discussion

In this study, we set out to determine whether established neural signatures of spatial navigation also applied to navigation of arbitrary representational spaces, in the form of episodic memory search. To do this, we had human neurosurgical patients perform a verbal free-recall task, and asked whether MTL theta power and connectivity correlated with pairwise distance in semantic and temporal representational spaces. Across all subjects, we found a reliable correlation between distances in a 1-D semantic space and theta activity, though this finding did not generalize to higher dimensions. Increases in theta power/connectivity were not readily associated with short temporal transitions, though the effect emerged when we filtered for list performance, and thereby ensured a dynamic range of temporal context reinstatement.

These results support the notion that theta oscillations are a common neural substrate of spatial navigation and episodic memory search, as predicted by the cognitive mapping theory of MTL function. Hippocampal theta has been associated with spatial navigation in rodents (see [52] for a review) and humans [22, 21, 23], and is seen during spontaneous retrieval of information encountered in a spatial layout [19]. Findings during non-spatial memory tasks have been decidedly more mixed, with many studies showing a decrease in MTL theta power during encoding and retrieval (e.g. [29, 30]). Here, we reconcile these results by using statistical contrasts of spatial or temporal clustering, instead of successful-versus-unsuccessful memory (e.g. the SME). This procedure inherently controls for nonspecific attentional or arousal states, since successful recalls occur in both conditions.

As predicted by models of MTL function [53], a positive association between human episodic memory and hippocampal theta becomes apparent when a more nuanced statistical contrast is employed. Not only does this harmonize results between the spatial and episodic memory literatures, it suggests the presence of a confound in the popular SME (or retrieval analogues): a non-specific task contrast between successful and unsuccessful memory operations includes significant brain activity unrelated to contextual memory processing, perhaps reflecting visual attention, executive control, or general task engagement. Unfortunately, these processes might cause broad decreases in theta power – for reasons still unknown – obscuring the theta rhythms that underlie the formation and retrieval of cognitive maps.

Notably, the methods used this manuscript involved no special techniques for extracting oscillatory and non-oscillatory components of the power or phase spectrum (e.g. [54, 55]). In effect, our clustering-based task contrast accomplishes the same thing, by controlling for the broadband spectral tilt that tends to reduce theta power and occurs primarily during successful memory states. However, we suspect that oscillation-detection procedures applied to this dataset could also reveal positive theta effects even in a nonspecific task contrast.

Several prior studies have addressed similar questions, but did not typically report strong hippocampal theta effects. Long, et al. (2015) examined neural activity during word encoding by comparing subsequently temporally clustered vs. not-clustered recalls, finding no increase in hippocampal theta power [56]. However, theta power was not significantly decreased between the two conditions either, unlike the strong theta decrease seen in the classic SME reported in that same study. This suggests a possible underlying increase in hippocampal theta that was obscured by residual non-contextual processes, which are especially strong during initial encoding. A related scalp EEG study by Long, et al. (2017) examined encoding-related activity for semantically-clustered retrievals, and reported a trend towards increases in low-theta power [57] – a finding that may have been significant with direct recordings of hippocampal LFPs.

In this manuscript, we found that theta power and connectivity were only significantly associated with the first principal word2vec dimension for each word list. Additional semantic dimensions were found to explain recall clustering at the behavioral level, but beyond the second dimension, the marginal explained variance fully accounts for this increase. In this sense, our behavioral and neurophysiological results align: one semantic dimension accounts for the majority of (1) recall clustering behavior and (2) MTL theta activity.

However, we suspect that these low-dimensional findings are partially a function of the list structure in our free recall task. Since PCA was performed on the 12 words in each list, high dimensions are likely to contain substantial noise, and therefore not correlate with hippocampal theta. Session-level PCA – which includes hundreds of words – can capture reasonable signal at higher dimensions but was actually less correlated with recall clustering (and theta activity) than the list-level PCA projections, at all dimensionalities. It is therefore possible that extraction of the first list-level PCA dimension is analogous to what the brain does during this episodic memory task; a human will consider only the words presented in each list, and try to find the single semantic feature axis that best captures their variability. This feature axis then becomes a dimension along which MTL theta reflects inter-item distances, just as it could also support navigation along a single dimension of a spatial layout.

The findings here show that the same theta signature of spatial navigation occurs during spontaneous episodic recall of word items. We therefore re-conceptualize free recall as navigation of a representational space with time and semantics as the most salient dimensions. The idea of a 2-D semantic-temporal space suggests that other signatures of 2-D spatial navigation, such as hexadirectional modulation of neural activity, may also be detectable during episodic recall. Two groups recently showed that theta power in the human entorhinal cortex is hexadirectionally modulated during virtual navigation [58, 59]. In the fMRI literature, it has been shown that hippocampal activity encodes for spatial [60] and abstract representational distances [61], and that hexadirectional modulation of the BOLD signal is observed during mental travel through abstract or imagined spaces [62, 63]. If free recall truly constitutes a form of navigation through a semantic-temporal space, might we observe hexadirectionally modulated theta during episodic retrieval?

Neuroscientists have long sought to understand how the medial temporal lobe simultaneously supports spatial navigation and episodic memory. This manuscript is another link in a growing chain of evidence that says memory and navigation are fundamentally supported by the same underlying cognitive process: the formation of associations between items, reflected as distances in arbitrary representational spaces. Until recently, spatial layouts have been the most obvious way to study these spaces. Here we demonstrated that temporal and semantic distances between word items were also captured by a classic hippocampal signature of navigation, lending strong support to the idea that the MTL performs domain-general cognitive mapping.

## 4 Methods

### Human subjects

For behavioral analyses, 189 adult patients with medication-resistant epilepsy underwent a surgical procedure to implant subdural platinum recording contacts on the cortical surface and within the brain parenchyma. Contacts were placed so as to best localize epileptic regions. Data reported were collected at 8 hospitals over 4 years (2015-2018): Thomas Jefferson University Hospital (Philadelphia, PA), University of Texas Southwestern Medical Center (Dallas, TX), Emory University Hospital (Atlanta, GA), Dartmouth-Hitchcock Medical Center (Lebanon, NH), Hospital of the University of Pennsylvania (Philadelphia, PA), Mayo Clinic (Rochester, MN), National Institutes of Health (Bethesda, MD), and Columbia University Hospital (New York, NY). Prior to data collection, our research protocol was approved by the Institutional Review Board at participating hospitals, and informed consent was obtained from each participant. For electrophysiological analyses, a subset of 96 patients with at least one contact placed in the MTL were used.

### Free-recall task

Each subject participated in a delayed free-recall task in which they studied a list of words with the intention to commit the items to memory. The task was performed at bedside on a laptop. The recall task consisted of three distinct phases: encoding, delay, and retrieval. During encoding, lists of 12 words were visually presented. Words were selected at random, without replacement, from a pool of high frequency English nouns (http://memory.psych.upenn.edu/WordPools). Word presentation lasted for a duration of 1600 ms, followed by a blank inter-sitmulus interval of 800 to 1200 ms. Before each list, subjects were given a 10-second countdown period during which they passively watch the screen as centrally-placed numbers count down from 10. Presentation of word lists was followed by a 20 second post-encoding delay, during which time subjects performed an arithmetic task during the delay in order to disrupt memory for end-of-list items. Math problems of the form A+B+C=?? were presented to the participant, with values of A, B, and C set to random single digit integers. After the delay, a row of asterisks, accompanied by a 60 Hz auditory tone, was presented for a duration of 300 ms to signal the start of the recall period. Subjects were instructed to recall as many words as possible from the most recent list, in any order, during the 30 second recall period. Vocal responses were digitally recorded and parsed offline using Penn TotalRecall (http://memory.psych.upenn.edu/TotalRecall). Subjects performed up to 25 recall lists in a single session (300 individual words).

### Intracranial EEG recordings

iEEG signal was recorded using depth electrodes (contacts spaced 2.2-10 mm apart) using recording systems at each clinical site. iEEG systems included DeltaMed XlTek (Natus), Grass Telefactor, Nihon-Kohden, and custom Medtronic EEG systems. Signals were sampled at 500, 1000, or 1600 Hz, depending on hardware restrictions and considerations of clinical application. Signals recorded at individual electrodes were first referenced to a common contact placed intracranially, on the scalp, or mastoid process. To eliminate potentially confounding large-scale artifacts and noise on the reference channel, we applied a bipolar re-reference for power analyses, and a common average reference of MTL contacts for connectivity analyses. Spectral analyses avoided the 55-65 Hz and 110-130 Hz ranges to mitigate contamination by line noise. As determined by a clinician, any contacts placed in epileptogenic tissue or exhibiting frequent inter-ictal spiking were excluded from all subsequent analyses.

### Anatomical localization

To localize contacts to the hippocampus or parahippocampal gyrus, hippocampal subfields and MTL cortices were automatically labeled in a pre-implant, T2-weighted MRI using the automatic segmentation of hippocampal subfields (ASHS) multi-atlas segmentation method [64]. Post-implant CT images were coregistered with presurgical T1 and T2 weighted structural scans with Advanced Normalization Tools [65]. MTL depth electrodes that were visible on CT scans were then localized within MTL subregions; parahippocampal cortex, perirhinal cortex, and entorhinal cortex were combined to form our PHG label, while CA1, CA3, dentate gyrus, and subiculum were combined to form our hippocampus label. Exposed recording contacts were approximately 1-2mm in diameter and 1-2.5mm in length; the smallest recording contacts used were 0.8mm in diameter and 1.4 mm in length.

### Semantic and temporal clustering analyses

In this study, we asked whether theta power during episodic memory retrieval correlated with distances between word items presented in a free-recall task. Distances were measured either as (1) semantic distance, computed from Euclidean distance in word2vec subspaces, or (2) temporal distances, computed from the serial position of the word in a sequentially-presented list.

The procedure to compute semantic distances is outlined in Fig. 1. The word2vec [36] representation of semantic values for each word came from publicly available vectors, trained on a large (approx. 100 billion words) text corpus from Google News (https://code.google.com/archive/p/word2vec/). These vectors represent each word as a point in a 300-dimensional space. To estimate semantic similarity between words, PCA was applied to the 12×300 matrix representing all of the words presented in a given list, to extract between one and 10 principal components. Similarity was taken as the Euclidean distance between points when projected onto the extracted PCA dimensions. (e.g. Fig. 1D-C). Alternatively, PCA was applied to the 300×300 matrix representing all 300 words encountered in a given experimental session (“Session PCA,” see Fig. 2B-C). Clustering and neurophysiological analyses were performed using inter-item distances measured in PCA-derived subspaces of varying dimensionality, by extracting different numbers of principal components.

Euclidean distances were computed between each successive pair of words as they were spoken during the free recall period for each list (e.g. Fig. 3A). Distance were converted to a semantic clustering score, reflecting the rank of the distance for reach recall transition, relative to all possible transitions. This was done by finding the percentile rank for the transition between the first pair of recalled words, relative to all possible transitions that could have been made among all words in the list. Next, the just-spoken word is removed as a possibility, and the procedure is repeated through all recall transitions. The result is a score between 0 and 1 for each recall transition, where 1 reflects the closest-possible transition, and 0 reflects the furthest-possible transition. Repeated words were excluded from all behavioral analyses.

In Fig. 2, each subject is assigned a Z-score that captures their overall degree of semantic recall clustering. For each list, a random subset of words is drawn, matched for the actual number recalled on that list, and the clustering scores are recomputed and averaged across lists. This procedure is repeated 250 times, creating a null distribution of clustering scores for each subject. The true average clustering score is compared to the null distribution to generate a Z-score for each subject. Higher Z-scores indicate a greater overall degree of semantic clustering.

As a benchmark, WordNet-derived distances were also used to measure semantic similarity. WordNet is a database enabling the assessment of word similarity via lexical relations [37]. Specifically, we used the Wu-Palmer measure of similarity to measure lexical distance between sequentially recalled words. Procedures were otherwise identical to those used for word2vec similarities.

Temporal clustering scores are computed exactly as semantic clustering scores, using the serial position of each word in the 12-item list instead of locations in word2vec-derived spaces. Therefore, a temporal clustering score of 1 indicates a subject recalled two words in sequence that occurred in sequence, while a score of 0 indicates a transition between words that were presented as far apart in time as possible. Z-scores are computed as above, except the null distribution is created by shuffling the true recall order for each list, instead of drawing a random subset of words.

To assess the degree to which clustering scores reflect increased explained variance in our PCA procedure – as opposed to recall behavior itself – we used linear regression to remove the effect of marginal explained variance (Fig. 2C). Specifically, for each subject, we took the first-order difference of their clustering versus PCA dimension curve, yielding a new curve that indicates the marginal increase in clustering for each added PCA dimension. We used linear regression to residualize these curves on explained variance for each PCA dimension, with residuals higher than zero indicating behavioral clustering in excess of what could be predicted by explained variance alone.

### Correlating spectral power with transition distance

To correlate spectral power with transition distances during episodic memory retrieval, for each subject we extracted 1-second intervals of iEEG from all (bipolar) electrodes placed in the MTL, immediately prior to each recall vocalization onset (Fig. 3B). Repeated words were excluded from all electrophysiological analyses. Spectral power was measured using the multitaper method as implemented in the MNE Python software package [66]. For each interval, theta power was taken as the averaged log-transformed power from 4-8 Hz, using a time-bandwith product of 4 and excluding tapers with <90% spectral concentration. Gamma power was taken as the average log power from 30-55 Hz, and high-frequency broadband was taken as the average from 70-150 Hz, with the 110-130 Hz range excluded to mitigate line noise artifact. Power was averaged across all electrodes in each region-of-interest (either hippocampus or PHG), and z-scored across all retrieval events.

As shown in Fig. 3C, z-scored power was Pearson correlated with semantic/temporal transition distances for all recalls in each experimental session, yielding a correlation coefficient for each subject and each session. Only lists with at least three recall transitions (i.e. four recalled words) were included, and any experimental session with fewer than 10 total recall transitions was excluded from all analyses. Correlation coefficients were Fisher z-transformed and averaged across sessions, yielding a final correlation score for each subject, region, and frequency band. Two-tailed 1-sample *t*-tests against 0 were used to determine whether, across subjects, power was significantly correlated with semantic or temporal transition distances (Fig. 4).

Because correlations do not indicate the actual degree of change in power, we also analyzed the association between power and transition distance by binning clustering scores into “short” (above 0.75) and “long” (below 0.25) categories, and taking the average power across all instances in each bin (see Fig. 3D and Fig. 6).

### Connectivity analyses

To analyze the relationship between intra-MTL connectivity and semantic/temporal transition distances during retrieval, we followed as similar approach as with spectral power. Connectivity was measured as the phase locking value (PLV), which reflects the consistency of phase differences between two electrodes either across trials or time [48]. In this study, we assessed PLV over time, which yielded a single-trial measure of connectivity between every pair of electrodes in the MTL for each retrieval event. To get a continuous measure of phase over time, we used the Morlet wavelet transform (5 cycles) with 1-second buffers to remove edge effects. PLVs were assessed for the theta and gamma bands separately, and averaged across all electrode pairs that spanned the hippocampus and PHG. We note that, to avoid analyzing connectivity between bipolar pairs which share a common monopolar contact, we used an MTL-average reference for connectivity analyses; i.e. subtracting the average signal across all channels placed anywhere in bilateral MTL. Procedures were otherwise identical to those described in “Correlating spectral power with transition distance” – hippocampal-PHG PLV was correlated with semantic/temporal transition distance across all retrieval events for each subject, generating a single connectivity-distance correlation measure for each subject. 56 subjects with contacts placed anywhere in the PHG and hippocampus were included in this analysis.

## Data and code availability

Raw electrophysiogical data used in this study is freely available at http://memory.psych.upenn.edu/Electrophysiological_Data. Analysis code and processed data is available on a public interactive Jupyter Note-book (https://notebooks.azure.com/esolo/projects/theta-semantics). Other custom processing scripts are available by request to the lead author.

## Acknowledgments

We thank Blackrock Microsystems and Medtronic for providing neural recording equipment. This work was supported by National Institutes of Health grant MH55687, as well as the DARPA Restoring Active Memory (RAM) program (Cooperative Agreement N66001-14-2-4032). We are indebted to all patients who have selflessly volunteered their time to participate in our study. The views, opinions, and/or findings contained in this material are those of the authors and should not be interpreted as representing the official views or policies of the Department of Defense or the U.S. Government. We also thank Dr. Nora Herweg for providing valuable feedback on this work. Icons in Fig. 1E by Freepik from www.flaticon.com.

