## Supplemental Figures for "Hippocampal theta codes for distances in semantic and temporal spaces"

### Supplemental Information (3 figures)

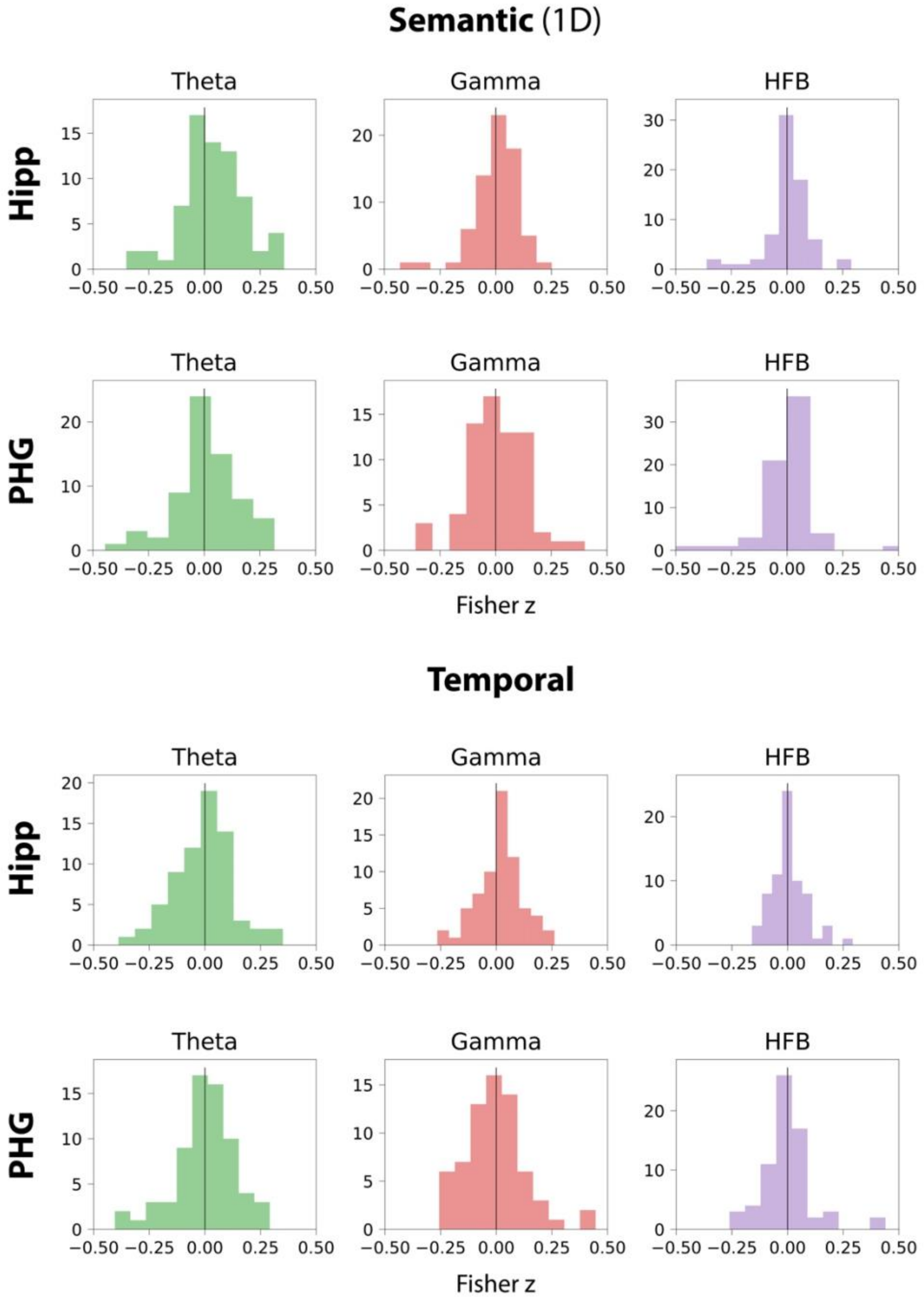


**Figure S1. Distribution of subject correlations** (Fisher z-transformed) for each region-feature-frequency combination. Data are summarized as bar plots in Figure 4. Theta (4-8 Hz), gamma (30-55 Hz), high-frequency broadband (HFB; 70-150 Hz). Only hippocampal theta (top left) was significantly predictive of semantic transition distances in 1-D spaces.

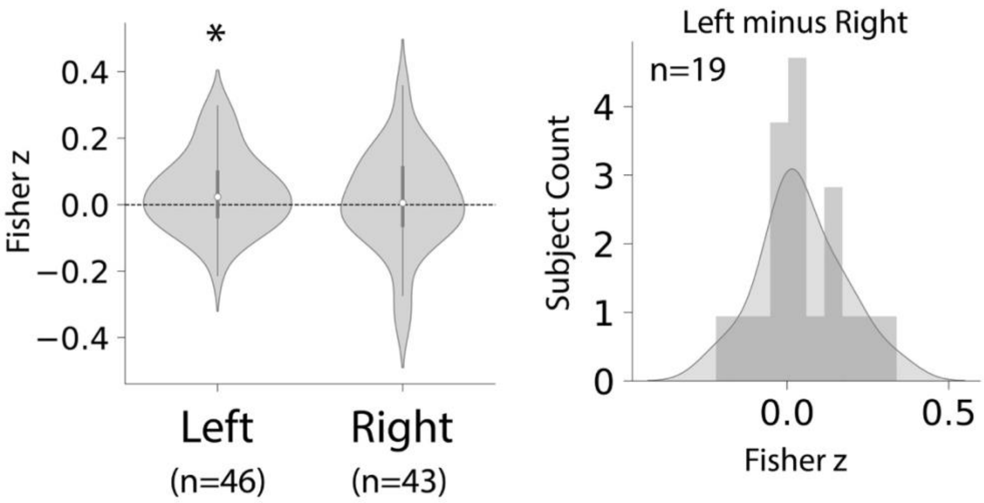


**Figure S2. Differences in theta power between left and right hippocampus.** *Left:* The correlation between hippocampal theta and semantic transition distance in (A) is driven primarily by activity in the left hippocampus; tested separately, left hippocampus (n=46 subjects) show a significant effect $t(45)=2.29$, $P=0.027$), unlike the right $t(42)=0.60$, $P=0.55$). *Right*: Testing for differences within subjects with bilateral hippocampal coverage, theta-distance correlations are higher in the left than the right, but not significantly so ($t(18)=1.49$, $P=0.15$).

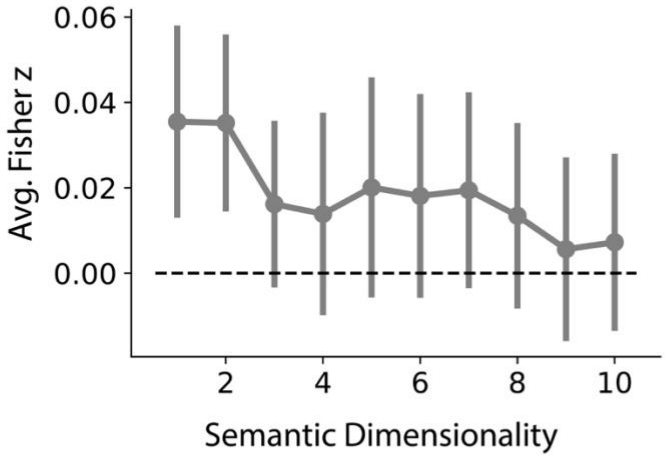


**Figure S3.** Population-level correlation between hippocampus-PHG theta coupling and semantic distance at varying dimensionalities, with MTL theta power regressed out. Same analysis as Figure 5E, though hippocampus-PHG phase locking was residualized on MTL theta power prior to correlation with semantic distance. One-dimension, $t$=1.58, $P$=0.12; two-dimensions, $t$=1.70, $P$=0.09.
